# BacTermFinder: A Comprehensive and General Bacterial Terminator Finder using a CNN Ensemble

**DOI:** 10.1101/2024.07.05.602086

**Authors:** Seyed Mohammad Amin Taheri Ghahfarokhi, Lourdes Peña-Castillo

## Abstract

A terminator is a DNA region that ends the transcription process. Currently, multiple computational tools are available for predicting bacterial terminators. However, these methods are specialized for certain bacteria or terminator type (i.e., intrinsic or factor-dependent). In this work, we developed BacTermFinder using an ensemble of Convolutional Neural Networks (CNNs) receiving as input four different representations of terminator sequences. To develop BacTermFinder, we collected roughly 41k bacterial terminators (intrinsic and factor-dependent) of 22 species with varying GC-content (from 28% to 71%) from published studies that used RNA-seq technologies. We evaluated BacTermFinder’s performance on terminators of five bacterial species (not used for training BacTermFinder) and two archaeal species. BacTermFinder’s performance was compared with that of four other bacterial terminator prediction tools. Based on our results, BacTermFinder outperforms all other four approaches in terms of average recall without increasing the number of false positives. Moreover, BacTermFinder identifies both types of terminators (intrinsic and factor-dependent) and generalizes to archaeal terminators. Additionally, we visualized the saliency map of the CNNs to gain insights on terminator motif per species. BacTermFinder is publicly available at https://github.com/BioinformaticsLabAtMUN/BacTermFinder.

## 1 Background

Transcription starts at the promoter region and ends at the terminator region. A terminator is a DNA segment that indicates the end of a gene or operon [55]. The termination of transcription is a process crucial for the accurate synthesis of RNA. In prokaryotes, termination can occur through either factor-dependent or factor-independent mechanisms, with the latter known as intrinsic termination. Intrinsic terminators are a DNA region rich in cytosine (C) and guanine (G) nucleotides followed by a polyuridine (U) sequence. The RNA created from the CG-rich region binds with itself, forming a hairpin structure which causes the RNA polymerase to stall. The weak base pairing between the adenine (A) nucleotides of the DNA template and the Us of the RNA transcript let the transcript to detach from the template thus, terminating the transcription [57]. While intrinsic termination requires only cis-acting elements on nascent RNAs, factor-dependent termination depends on both cis-acting RNA elements and a protein such as Rho (*ρ*) which mediates termination via three distinct mechanisms [62].

Identifying terminators is crucial for understanding operon structure and transcriptional regulation. As both terminator types rely, at least partially, on cis-acting elements on the DNA template, they can be identified from genomic sequence. This can be done via computational identification. Genome-wide terminator identification via computational approaches is more affordable and efficient than via wet lab approaches. Since the development of TransTermHP [36], which is arguably the most cited terminator prediction tool, there have been several computational tools developed for bacterial terminator prediction (summarized in Table 1). However, as one can see in Table 1, these tools have either 1) focused on few (one to three) bacterial species (mostly *Escherichia coli* and *Bacillus subtilis*), 2) used a relatively small number of terminators for generating their model, 3) predicted a single terminator type (factor-dependent or intrinsic), or 4) a combination of the above. Thus, there is still the need for a species- and terminator type-independent tool for predicting bacterial terminators. The availability of genome-wide transcription termination sites (TTSs) identified by RNA-seq technologies such as Term-Seq [14], Send-seq [33], SMRT-cappable [69], RendSeq [38], RNATag-seq [60], and dRNA-seq [59] in several bacterial species opens the door to generate a species-agnostic machine learning-based model using a large number (i.e., thousands) of terminator sequences of a wide range of bacterial species. Here, we generated such a method by 1) gathering a large collection of TTSs from published studies, 2) exploring thousands of features to represent (encode) terminator sequences, 3) generating and assessing eleven different machine learning models to identify bacterial terminators, and 4) comparatively assessing the performance of our best model (BacTermFinder) with the performance of four other bacterial terminator prediction methods (namely, TermNN [8], iTerm-PseKNC [19], RhoTermPredict [16] and TransTermHP [36]). Our results show that BacTermFinder can detect intrinsic and factor-dependent terminators and even archaeal terminators at a higher recall rate than current tools.

**Table 1:** Summary of tools for predicting bacterial terminators. # of terminators indicates how many terminator sequences were used in the study. The # of species indicates how many bacterial species were considered. CM is covariance model, DA = discriminant analysis, DL = deep learning, DP = dynamic programming, and ML = machine learning.

| Methods | Year | Terminator type | Method | Software available? | # of terminators | # of species |
| --- | --- | --- | --- | --- | --- | --- |
| TermNN [8] | 2022 | Intrinsic | DL | Yes | 1175 | 2 |
| ITT pred [26] | 2021 | Intrinsic | Statistical | No | 137 | 1 |
| iterb-PPse [18] | 2020 | Both | ML | No | 928 | 2 |
| iTerm-PseKNC [19] | 2019 | Both | ML | Yes | 852 | 1 |
| RhoTermPredict [16] | 2019 | Factor-dep. | DP | Yes | 1298 | 3 |
| OPLS-DA [52] | 2018 | Factor-dep. | DA | Yes | 104 | 2 |
| PASIFIC [51] | 2017 | Intrinsic | ML | Yes | 330 | 89 |
| RNIE [23] | 2011 | Intrinsic | CM | Yes | 1062 | 2 |
| TransTermHP [36] | 2007 | Intrinsic | DP | Yes | N/A | N/A |

## 2 Methods

### 2.1 Terminator sequences collection

We searched the NCBI PubMed [1] and GEO database [2] using as keywords Term-Seq, Send-seq, SMRT-cappable, RendSeq, RNATag-seq, and dRNA-seq to find published studies which identified bacterial TTSs. For each identified study, using an IPython Notebook [37] and pandas data manipulation library [53], we stored in a BED file the genomic location of identified TTSs (provided in the supplementary material of the corresponding publication). With BEDTools’ [54] slopeBed and FastaFromBed commands, we extracted the genomic sequences corresponding to 100 nucleotides (nts) flanking the TTSs (50 nts on either side) into a FASTA file. In each operation, strandedness was taken into account. We chose 50 nts on either side of the TTSs as a previous study [44] found intrinsic terminator regions in *E. coli* as long as 65 nts. Thus, a sequence of 100 nts long would be enough to contain this region and allow for some variation. Terminators present in plasmids were disregarded. We also collected archaeal TTSs during this process and kept these for the comparative assessment. Additionally, we collected TTSs available in the following databases: RegulonDB [22], DBTBS [30], and BSGatlasDB [24]. As we collected multiple data sets for some of the bacterial species, we needed to remove duplicated sequences from our data. To achieve this, we merged genomic locations if they had at least a 60% genomic location overlap. When the merged sequence was bigger than 100 nts, we symmetrically removed the extra nts from the beginning and the end of the sequence. Additionally, sequences containing ambiguous characters (such as N, Y, etc) were removed. As some of the studies we collected data from have used different genome assemblies for the same bacterium, after getting the terminator sequences, we removed duplicated sequences (i.e., sequences 100% identical). The bacterial species with their number of non-redundant terminators used for training and comparative assessment are listed in Tables 2 and 3, respectively. Details about each study are listed in Supplementary Table 1.

**Table 2:** Number of non-redundant terminators per genome accession used for training. There is a total of 36,542 terminator sequences.

| Data source | Species | Genome Accession | Terminators | GC (%) |
| --- | --- | --- | --- | --- |
| [14, 13] | *Bacillus subtilis* 168 | AL009126.3 | 1800 | 42.90 |
| [63, 49, 39, 24, 32, 27] | *Bacillus subtilis* 168 | NC_000964.3 | 4872 | 42.90 |
| [39] | *Caulobacter crescentus* NA1000 | CP001340.1 | 341 | 66.22 |
| [21] | *Clostridioides difficile* 630 | CP010905.2 | 1646 | 28.63 |
| [20] | *Dickeya dadantii* 3937 | NC_014500.1 | 1786 | 55.50 |
| [15] | *Escherichia coli* BW25113 | CP009273.1 | 1095 | 50.06 |
| [39, 32, 33, 58, 69, 3, 12] | *Escherichia coli* str. K-12 substr. MG1655 | NC_000913.3 | 4139 | 50.07 |
| [64] | *Pseudomonas aeruginosa* PAO1 | NC_002516.2 | 805 | 65.61 |
| [50] | *Staphylococcus aureus* JKD6009 | LR027876.1 | 978 | 32.41 |
| [5] | *Staphylococcus aureus* NCTC 8325 | NC_007795.1 | 566 | 32.40 |
| [61] | *Streptococcus pneumoniae* D39V | CP027540.1 | 747 | 39.14 |
| [66] | *Streptococcus pneumoniae* TIGR4 | NC_003028.3 | 1810 | 39.13 |
| [41, 42] | *Streptomyces avermitilis* MA-4680 | BA000030.4 | 2006 | 69.72 |
| [28, 42] | *Streptomyces clavuligerus* ATCC27064 | CP027858.1 | 1583 | 71.64 |
| [42] | *Streptomyces coelicolor* M145 | NC_003888.3 | 1308 | 71.10 |
| [42, 29] | *Streptomyces griseus* NBRC13350 | NC_010572.1 | 2724 | 71.21 |
| [43, 42] | *Streptomyces lividans* TK24 | CP009124.1 | 1999 | 71.22 |
| [42] | *Streptomyces tsukubaensis* NBRC108819 | CP020700.1 | 1283 | 70.85 |
| [39] | *Vibrio natriegens* ATCC 14048 | CP009977.1 | 905 | 44.66 |
| [39] | *Vibrio natriegens* ATCC 14048 | CP009978.1 | 257 | 44.10 |
| [34] | *Vibrio parahaemolyticus* RIMD 2210633 | NC_004603.1 | 1852 | 44.74 |
| [65] | *Zymomonas mobilis* ZM4 = ATCC 31821 | CP023715.1 | 2040 | 45.67 |

**Table 3:** Number of non-redundant terminators per species used for comparative assessment for a total of 4,626 and 3,581 bacterial and archaeal terminators, respectively.

| Data source | Species | Genome Accession | Terminators | % GC | Seq. Tech. |
| --- | --- | --- | --- | --- | --- |
| **Bacteria** |  |  |  |  |  |
| [17] | *Mycobacterium tuberculosis* H37Rv | AL123456.3 | 2202 | 64.69 | Term-Seq |
| [56] | *Streptococcus agalactiae* NEM316 | NC_004368.1 | 655 | 35.12 | dRNA-Seq |
| [42] | *Streptomyces gardneri* ATCC 15439 | CP059991.1 | 870 | 70.73 | Term-Seq |
| [11] | *Synechocystis* PCC 6803 | NC_000911.1 | 553 | 47.04 | Term-Seq |
| [31] | *Synechocystis* PCC 7338 | CP054306.1 | 346 | 47.14 | Term-Seq |
| **Archaea** |  |  |  |  |  |
| [6] | *Haloferax volcanii* DS2 | NC_013967 | 1227 | 65.69 | Term-Seq |
| [45] | *Methanococcus maripaludis* S2 | NC_005791 | 2354 | 32.63 | Term-Seq |

### 2.2 Confirming the sequence length for terminator identification

To confirm that a sequence pattern was within the 100 nts extracted, we used relative nucleotide frequency graphs to visualize whether a distinct pattern was present within this region of interest (ROI). The relative log nucleotide frequency for a specific position was calculated by dividing the total count of each nucleotide in that position by the number of terminators in that set and then applying a log base 2 function. The log2 ratio of the relative nucleotide frequency in the ROI vs random genomic regions for each position and nucleotide was obtained by subtracting from the relative log nucleotide frequency in the ROI the relative log nucleotide frequency in randomly selected genomic sequences of the same length.

### 2.3 Non-terminator sequences generation

To train machine learning-based models, we needed instances of non-terminators of the same length and from the same genome as the terminator-containing sequences. To generate these negative examples, we used BEDTools’ shuffle command to randomly generate genomic coordinates different from the terminator regions. We allowed a maximum sequence overlap between positive and negative sequences of 20 nts. A ratio of 1-10 positive to negative was used as it was shown in [10] that there should be more negative than positive instances to lower the false-positive rate during genome scan. Furthermore, the terminator detection problem has a natural imbalance between terminator and non-terminator sequences, as the estimated number of terminators in a bacterial genome is relatively tiny compared to the number of all possible non-terminator sequences of the same length. Henceforth, we refer to our complete data set (including terminator and non-terminator sequences) as BacTermData.

### 2.4 Feature engineering and selection

BacTermData contains DNA sequences consisting of ATCG characters. However, machine learning methods expect to receive a numerical representation of the sequences as input. There are many approaches to numerically representing sequences, such as one-hot (binary) encoding, k-mer frequencies, etc. As it is not feasible to determine a priori which representation would generate the “best-performing” machine learning model, one needs to try out several distinct representations. Here, the best-performing model refers to a model that maximizes a specific performance metric, such as the F1-score or area under the precision-recall curve (AUPRC). There are several libraries or software (e.g., MathFeature [7], iLearnPlus [9], RepDNA [46]) to generate features from DNA sequences. Here we decided to use iLearn-Plus to represent the sequences in BacTermData, as it can generate a wide variety of feature sets. Using iLearnPlus, we computed 6208 features (belonging to 28 feature sets) per sequence. A feature set contains features created by one feature generation method (e.g., one-hot encoding). After generating the features, we applied feature selection methods to identify informative features.

To measure the importance of the features, we used two methods: SHAP [47] and Gini measure of Light Gradient Boosting Machine (LGBM) [35]. We used an iterative algorithm to drop features: 1) train an LGBM model with all remaining features and calculate feature importance values using SHAP and Gini measure, 2) remove features in the bottom 20% after sorting the features based on their importance as per both feature importance methods, 3) repeat step 1 and 2 until the average of the AUPRC obtained during 10-fold cross-validation decreases. We observed a decrease in the AUPRC after reaching 1694 features, down from 6208 features (Fig. 1). We called these 1694 features (belonging to 22 feature sets) the 22-sets features. Supplementary Table 2 describes the 1694 selected features.

**Figure 1:**
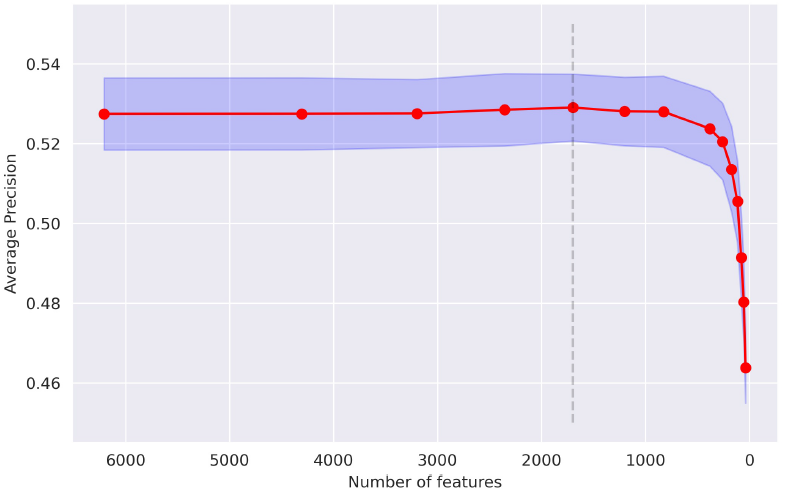
Average precision (an estimate of AUPRC) as a function of the number of features included in the training set. Each dot indicates an iteration of the algorithm used to remove non-informative features. The shaded area is the standard deviation of 10 cross-validation folds. The vertical dash line indicates the point where a drop of AUPRC occurred.

Generating the 1694 features with ILearnPlus is computationally intensive (roughly, it processes 30 sequences/second in a High-Performance Computing Cluster (HPC cluster)). To reduce the computational requirements, we selected the complete six feature sets with the most important features. These six features sets comprised 60% of the 1694 features (Supplementary Table 2) and called the corresponding features the 6-sets features. We also selected individual feature sets such as ENAC and one-hot (OH) encoding which preserve the spatial location of patterns in the ROI.

### 2.5 Machine learning approaches

We utilized convolutional neural networks (CNNs) and fully-connected neural networks (FCNNs) as they have been shown to perform well in various domains [40]. We also used a boosting method because of boosting methods’ ability to handle tabular data better than other techniques [25]. Table 4 describes the ML approaches considered. Table 5 describes the CNNs architecture.

**Table 4:** Machine learning approaches considered. Feature sets are described in Supplementary Table 2.

| # | ML Method | Features | Description |
| --- | --- | --- | --- |
| 1 | CNN | PS2, ENAC, OH, 6-sets features | Fusion<sup>1</sup> |
| 2 | CNN | OH, 6-sets features | Fusion |
| 3 | CNN | PS2, ENAC, OH, 6-sets features | Append<sup>2</sup> |
| 4 | CNN | PS2 | Single<sup>3</sup> |
| 5 | CNN | OH | Single |
| 6 | CNN | ENAC | Single |
| 7 | CNN | NCP | Single |
| 8 | CNN | PS2, ENAC, OH, NCP | Ensemble<sup>4</sup> |
| 9 | FCNN | 6-sets features | FCNN trained on 6-sets features |
| 10 | LGBM | 6-sets features | A LGBM trained on 6-sets features |
| 11 | LGBM | 22-sets features | A LGBM trained on 22-sets features |

Train one CNN for each of the listed feature sets followed by fully-connected layer(s).
Train a CNN receiving as input the concatenated listed feature sets followed by fully-connected layer(s).
Train a CNN receiving as input the listed feature set followed by fully-connected layer(s).
Train independent CNNs followed by fully-connected layer(s) on each listed feature set and the combined prediction is their average output.

**Table 5:** CNN architecture description.

| Layer Type | Hyperparameter | Value |
| --- | --- | --- |
| Conv1D | filters | 64 |
|  | kernel_size | 10 |
|  | activation | PReLU |
| AveragePooling1D | pool_size | 2 |
| Conv1D | filters | 64 |
|  | kernel_size | 10 |
|  | activation | PReLU |
| AveragePooling1D | pool_size | 2 |
| BatchNormalization | - | - |
| Conv1D | filters | 64 |
|  | kernel_size | 10 |
|  | activation | PReLU |
| Dropout | rate | 0.1 |
| Flatten | - | - |
| Dense (Dropout) | units & (rate) | 500 (0.3) |
|  | activation | PReLU |
| Dense (Dropout) | units & (rate) | 600 (0.3) |
|  | activation | PReLU |
| Dense (Dropout) | units & (rate) | 600 (0.3) |
|  | activation | PReLU |
| Dense (Dropout) | units & (rate) | 200 (0.4) |
|  | activation | PReLU |
| Dense (Dropout) | units & (rate) | 200 (0.4) |
|  | activation | PReLU |
| Dense | units | 1 |
|  | activation | Sigmoid |

The architecture of the FCNN is a dense layer containing 400 units and, using the ReLU activation function, is applied to the categorical input. This is followed by a dropout layer with a rate of 0.3 to reduce overfitting. This pattern of a dense layer with 400 units and ReLU activation followed by a dropout layer with a rate of 0.3 is repeated 6 times more. The output from these repeated layers is then fed into a dense layer with 200 units and parametric ReLU activation, followed by a dropout layer with a rate of 0.4. This configuration is repeated once more before the final output layer, which consists of a single unit with a sigmoid activation function to provide a binary classification output.

### 2.6 Model generation and assessment

We used stratified Monte Carlo cross-validation (SMCCV) [68] (also called repeated random subsampling) to select the optimal hyperparameters of the neural network models. Monte Carlo cross-validation is a method that randomly determines the training and validation set in different iterations. The positive-to-negative data ratio is maintained in each iteration as it is stratified. We used SMCCV with 100 folds and 10 iterations to find the optimal hyper-parameters for the models. With 100 folds, 99 % of the data is for training and 1% for testing on each fold. Neural networks were implemented in TensorFlow version 2.8.0. We used randomized cross-validation with 50 iterations to find the best hyper-parameters for the LGBM model (Table 6). We used the Python LGBM implementation version 3.3.3.

**Table 6:** Best hyperparameters for LightGBM Model.

| Hyperparameter | Value |
| --- | --- |
| boosting_type | goss |
| colsample_bytree | 0.8 |
| learning_rate | 0.1 |
| max_depth | 7 |
| min_child_weight | 1 |
| n_estimators | 3000 |
| num_leaves | 64 |
| objective | cross_entropy_lambda |
| random_state | 42 |
| reg_lambda | 30 |
| unbalance | True |
| n_jobs | -1 |

We used SMCCV to assess the models’ performance, and calculated average precision, recall, and F-score per species. We calculated score thresholds for classification that maximize the F0.5, F1, and F2 scores. Average precision is calculated as AP =∑_*n*_ *(R*_*n*_*− R*_*n −* 1_)*P*_*n*_ where *R*_*n*_ and *P*_*n*_ are the precision and recall at the *n*^th^ threshold. Recall is the proportion of actual positive instances correctly identified by a model 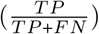, while precision measures the proportion of predicted positive instances that are actually true positives 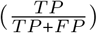 where *TP* is the number of true positives, *FN* is the number of false negatives, and *FP* is the number of false positives. F-score is calculated as F-score 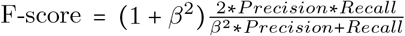 where *β* can be 0.5, 1, or 2 for the corresponding F Score.

## 3 Results and Discussion

### 3.1 Clearly visible pattern in ROI in BacTermData

We visualized the log2 ratio of the relative nucleotide frequency across the ROI for each study data included in BacTermData and confirmed the presence of a clear pattern around the middle of the ROI in all the data. Fig. 2 shows the aggregated pattern around the TTS (middle of the sequence) in the terminator sequences in BacTermData. This indicates that there is indeed a pattern in BacTermData, which machine learning approaches could learn to identify terminator sequences. BacTermData consists of 41,168 bacterial terminator sequences (36,542 used for training) of 25 bacterial strains belonging to four phyla: Pseudomonadota (formerly Proteobacteria), Bacillota (formerly Firmicutes), Actinomycetota (formerly Actinobacteria) and Cyanobacteriota. To test the generalization capabilities of our final model, the terminator sequences of five bacterial strains (Table 3), including the only two belonging to the phylum Cyanobacteriota, were left out of training and used only for the comparative assessment.

**Figure 2:**
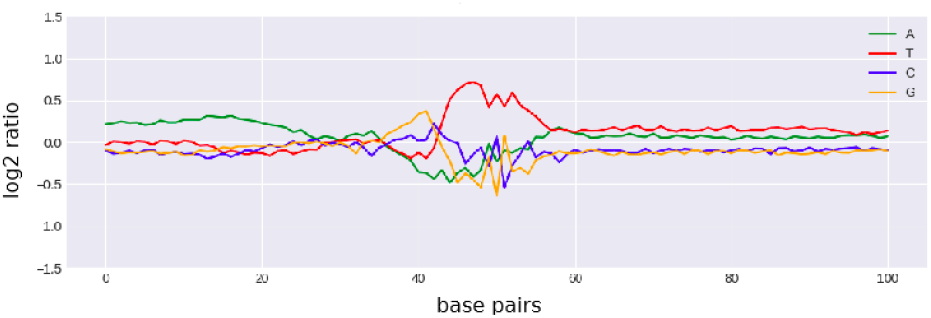
Log2 ratio of the relative nucleotide frequency of all terminator sequences in BacTermData.

### 3.2 Selecting a best performing model: BacTermFinder

We assessed various ML approaches and sequence representation combinations (Table 4). This assessment involved training and evaluating these combinations on the data shown in Table 2 using Stratified Monte-Carlo Cross Validation (SMCCV). The SMCCV results are presented in Table 7. The LGBM model trained on the 22-sets features achieved performance comparable to that of single CNNs, however computing the 22-sets features is resource-intensive and time-consuming. During training the models, we noticed that the single CNN models were performing well based on average precision and thus, we decided to build an ensemble with these four models by simply averaging their outputs. This approach consisting of an ensemble of the four single CNNs (Fig. 3) achieved an average precision of 0.7080 ± 0.0248, out-performing all the other approaches considered (Table 7). We selected this as our final model and called it BacTermFinder. Fig. 4 shows the Precision-Recall curve (PRC) and Receiver Operating Characteristic curve (ROC) of BacTermFinder over all SMCCV iterations. Clearly, BacTermFinder’s performance is well above a random classifier’s performance (dashed lines in Fig. 4).

**Table 7:** SMCCV average precision ± standard deviation. The highest SMCCV average precision is highlighted in bold. See Table 4 for a description of the ML approaches.

| # | Approach | Average Precision ± s.d. |
| --- | --- | --- |
| 1 | CNN_Fusion | 0.5955 ± 0.0232 |
| 2 | CNN_Fusion | 0.6553 ± 0.0252 |
| 3 | CNN_Append | 0.5949 ± 0.0257 |
| 4 | CNN_Single | 0.6738 ± 0.0253 |
| 5 | CNN_Single | 0.6775 ± 0.0214 |
| 6 | CNN_Single | 0.6544 ± 0.0284 |
| 7 | CNN_Single | 0.6730 ± 0.0243 |
| 8 | CNN_Ensemble | **0.7080 ± 0.0248** |
| 9 | FCNN | 0.5012 ± 0.0319 |
| 10 | LGBM | 0.5933 ± 0.0269 |
| 11 | LGBM | 0.6476 ± 0.0217 |

**Figure 3:**
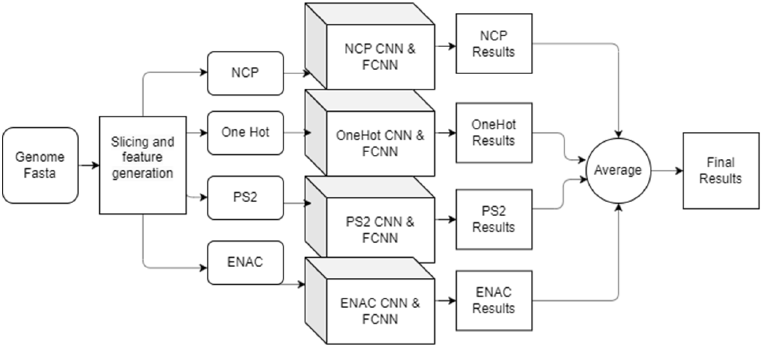
Depiction of BacTermFinder architecture.

**Figure 4:**
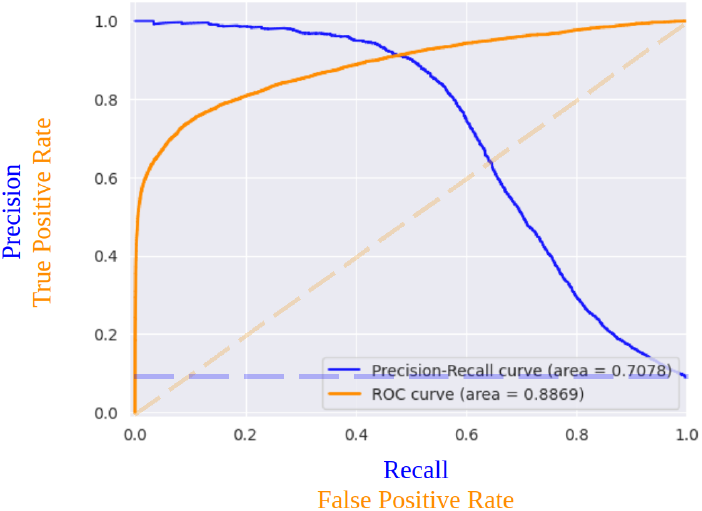
PRC in blue and ROC in orange of Bac-TermFinder on the aggregated results of SMCCV iterations. Dashed lines show the performance of a random classifier.

### 3.3 Effect of GC content on terminator identification

To look further into the variation in performance across bacterial species, we visualized the average precision per bacterium vs their GC content and coloured it by the phylum to see if there was any pattern (Fig. 5). A linear regression line fitted to this data indicates a relationship between GC content and average precision: As the GC content increases, the performance decreases (Spearman correlation value of -0.46, p-value 0.031). For a 10% increase in GC content, the average precision decreases by 0.038. Thus, BacTermFinder tends to achieve higher average precision in bacteria with lower GC content. Bacteria with high GC content tend to have more factor-dependent terminators [17]. Factor-dependent terminators do not always have the hairpin structure of intrinsic terminators [67], and thus, their sequence motif might be weaker. Additionally, the Rho utilization (rut) site is 60-90 nts upstream of the terminator [4], and thus, outside our ROI. These two issues might explain why BacTermFinder’s performance is lower on high GC bacteria. BacTermFinder’s performance is more consistent (with less variation) for Bacilota and Actinomycetota than for Pseudomonadota (Fig. 5). Further investigation is needed to understand the reasons for this.

**Figure 5:**
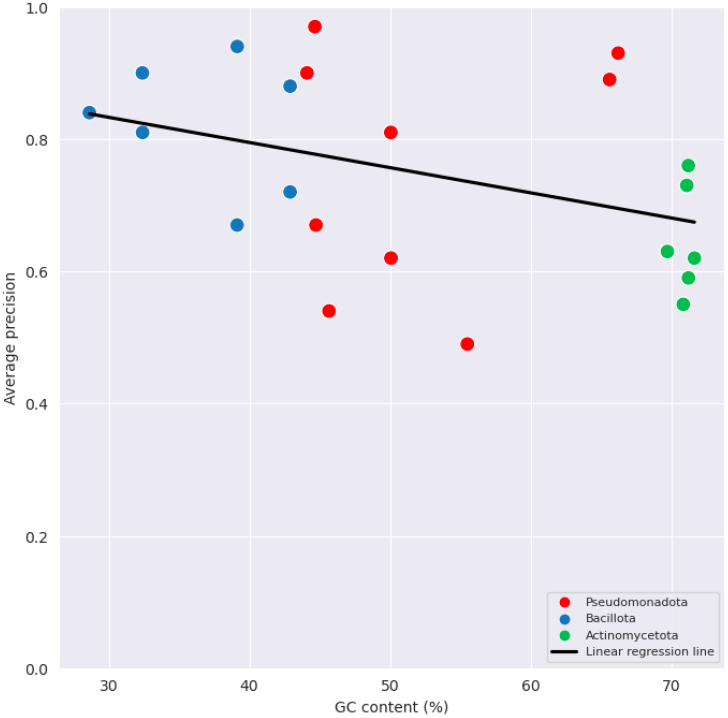
Average precision vs GC content of Bac-TermFinder SMCCV results per bacterial strain. Bacterial strains are coloured based on their phylum.

### 3.4 Assessing BacTermFinder’s predictions on independent validation data

We tested BacTermFinder on an independent set of bacterial and archaeal terminators not included in the training data (Table 3). For this assessment, we used the confidence threshold for classification that maximizes the F2 score. We visually looked at whether predicted terminators have a log2 ratio of the relative nucleotide frequency similar to that of experimentally identified terminators. As it can be seen from Fig. 6 for *S. agalactiae*, the log2 ratio of the relative nucleotide frequency of predicted terminators is similar to that of experimentally identified terminators. The line smoothness of Fig.6(B) is due to being a larger number of predicted terminators (12,813) vs experimentally identified terminators (655). The saliency map (Fig. 6(C)) shows the contribution of each nucleotide per position in BacTermFinder’s predictions. Negative numbers reduce the probability of predicting a terminator, while positive numbers increase the probability of predicting a terminator. Figs. 7 and 8 show the same visual analysis for the archaea *M. maripaludis* and the cyanobacterium *Synechocystis* PCC 6803, suggesting that BacTermFinder predicted terminators have similar sequence patterns to those present in experimentally determined terminators and that BacTermFinder is also suitable for finding archaeal terminators.

**Figure 6:**
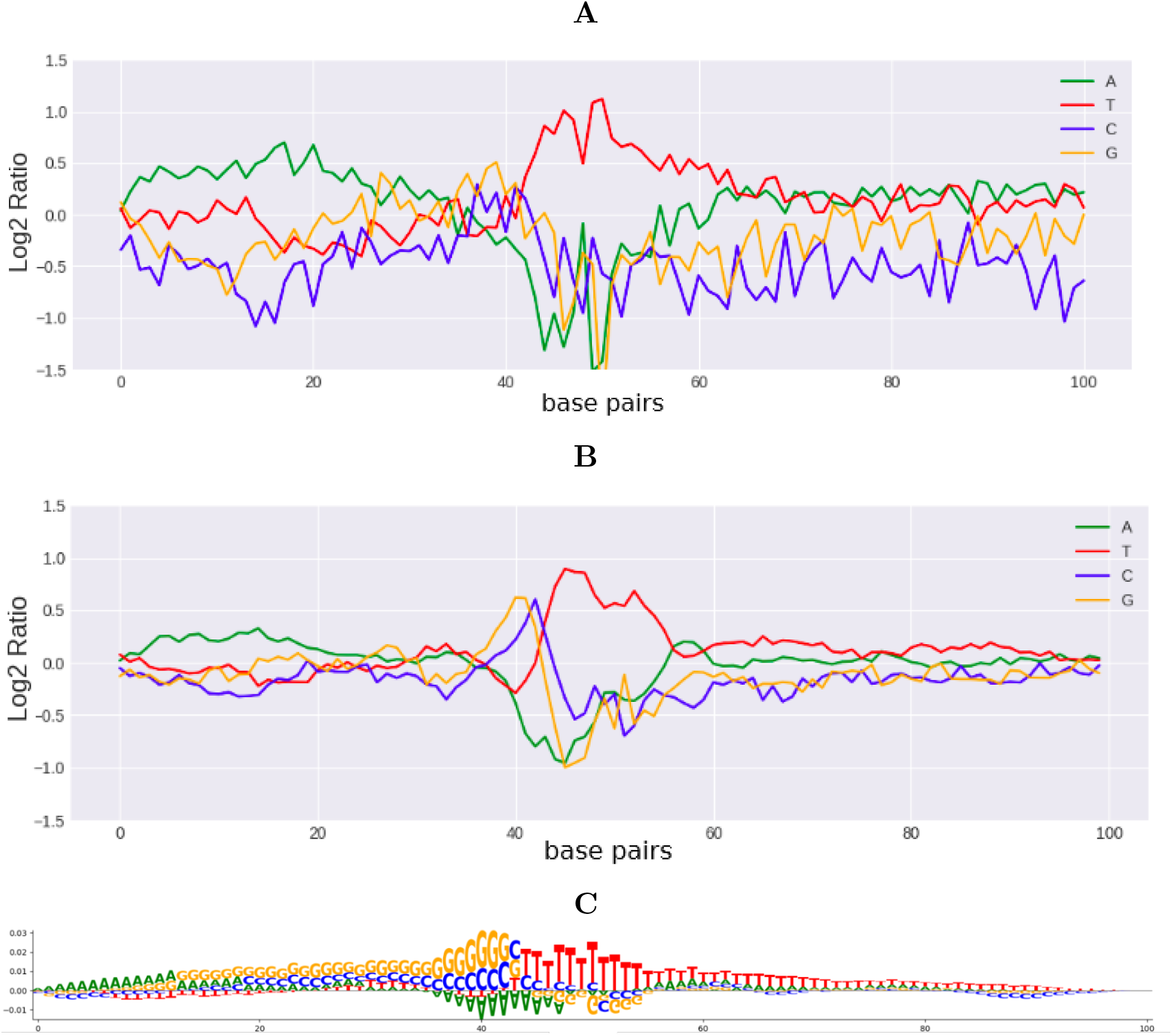
Visualization of *S. agalactiae* experimentally identified terminators (A) vs BacTermFinder genome-wide predictions (B). **A**. Log2 ratio of the relative nucleotide frequency of experimentally determined terminators. **B**. Log2 ratio of the relative nucleotide frequency of genome-wide predicted terminators. **C**. Saliency map generated using using [48].

**Figure 7:**
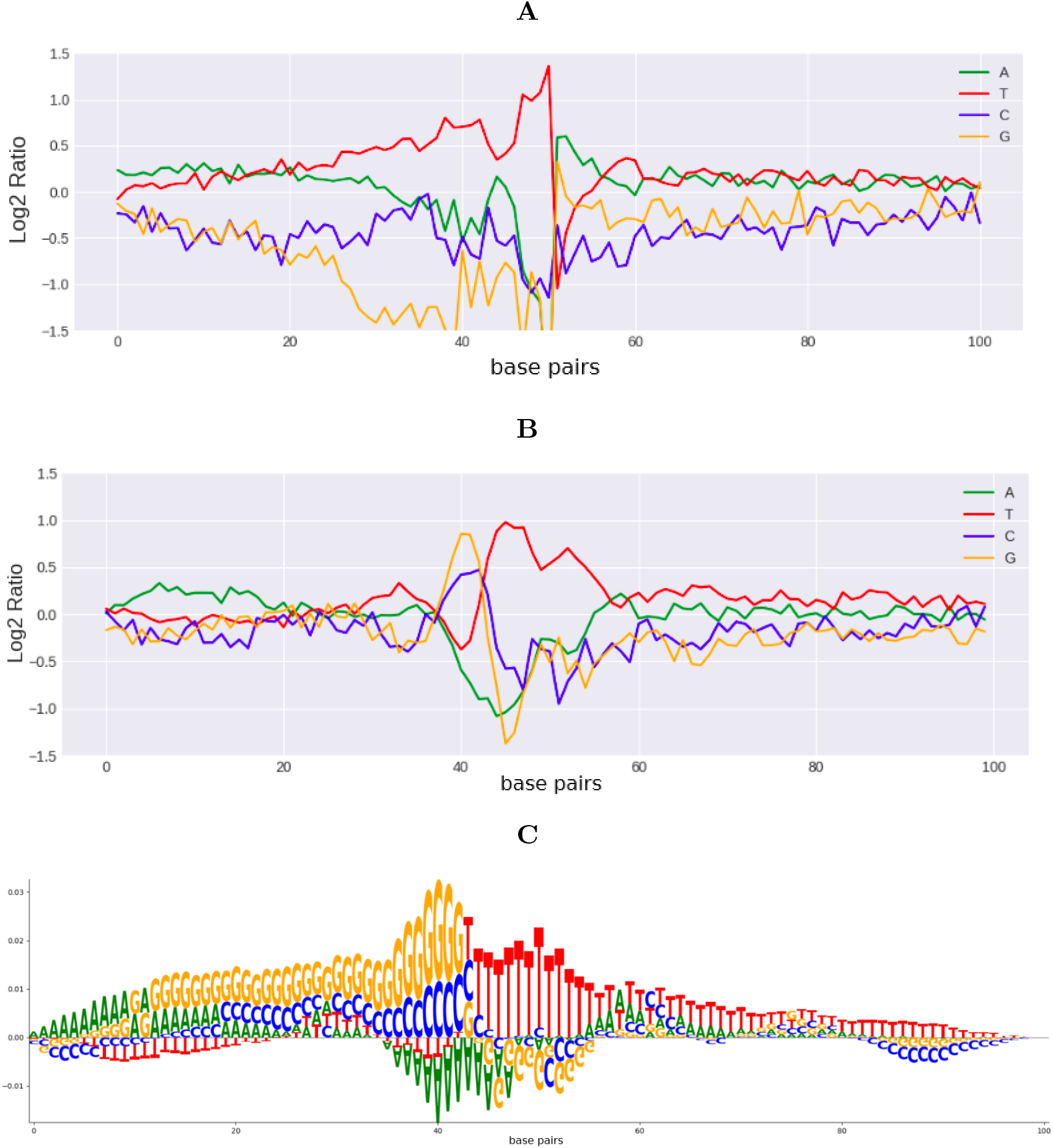
Visualization of *M. maripaludis* S2 experimentally identified terminators (A) vs BacTermFinder genome-wide predictions (B). **A**. Log2 ratio of the relative nucleotide frequency of experimentally determined terminators. **B**. Log2 ratio of the relative nucleotide frequency of genome-wide predicted terminators. **C**. Saliency map generated using [48].

**Figure 8:**
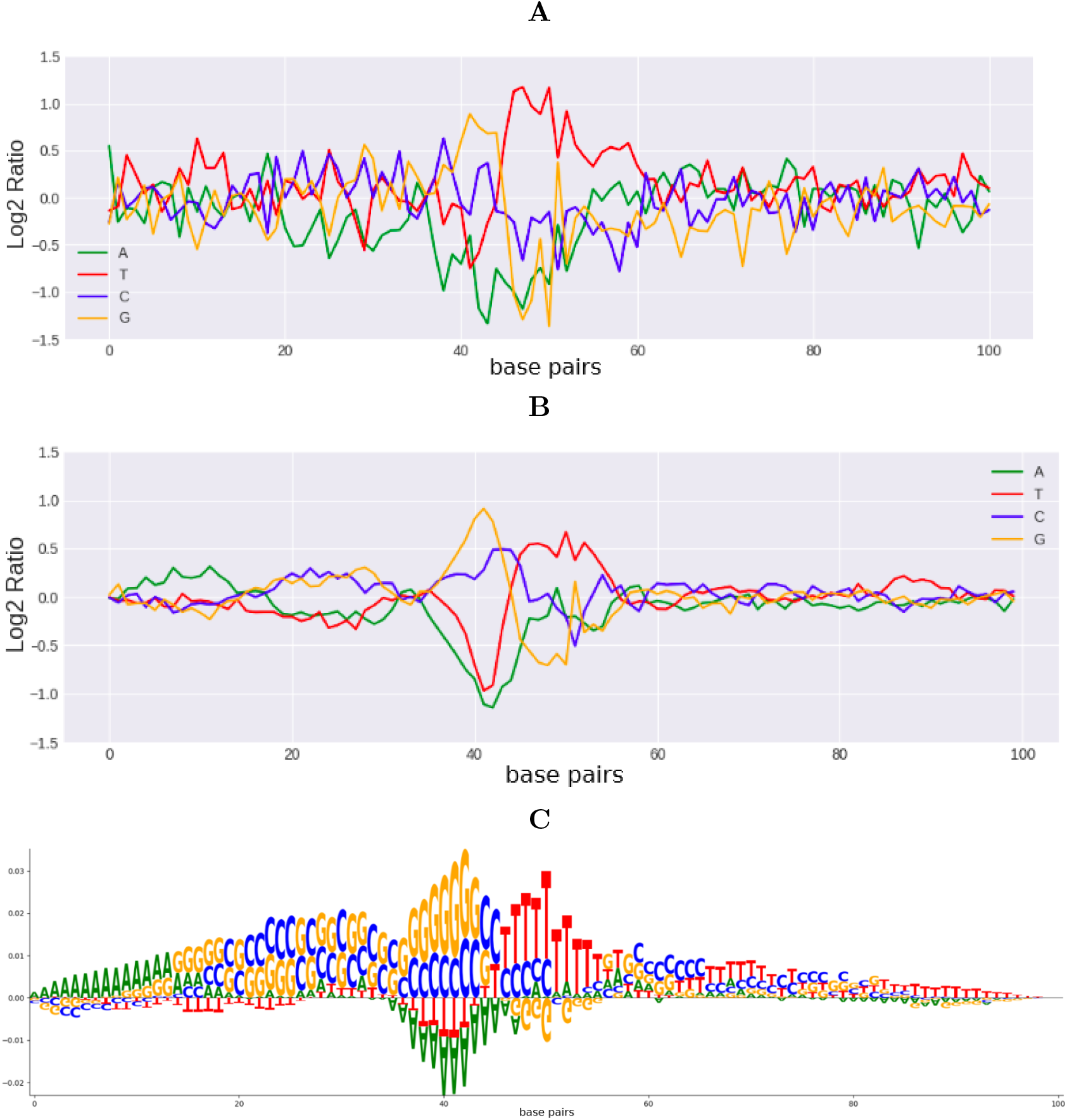
Visualization of *Synechocystis* PCC 6803 experimentally identified terminators (A) vs BacTermFinder genome-wide predictions (B). **A**. Log2 ratio of the relative nucleotide frequency of experimentally determined terminators. **B**. Log2 ratio of the relative nucleotide frequency of genome-wide predicted terminators. **C**. Saliency map generated using [48].

We noticed some commonalities in the terminator sequence patterns shown in the saliency maps of *H. volcanii, M. tuberculosis*, and *S. gardneri* (Supplementary Figs. 1-3(C)) which are all organisms with high GC content in their genome. Similarly, the terminator sequence patterns of *S. agalactiae* and *M. maripaludis* as per their saliency maps (Figs. 6(C) and 7(C)) show an increasing preference for Gs and Cs peaking around position 43 followed by a long poly-T. These two organisms have some of the lowest GC content in our data. Finally, the two Synechocystis have their own distinct shared pattern (Fig. 8(C) and Supplementary Fig. 4(C)).

### 3.5 Comparative assessment for genome-wide terminator prediction

We compared BacTermFinder’s performance with that of TermNN [8], iTerm-PseKNC [19], RhoTermPredict [16] and TransTermHP [36] on terminators of five bacterial species (Table 3) not used for generating our model. We decided to include TermNN, RhoTermPredict and iTerm-PseKNC in the comparative assessment because they are the most recently developed tools available for predicting intrinsic, factor-dependent and both types of terminators, respectively (Table 1). We included TransTermHP because, as mentioned earlier, is the most cited tool for bacterial terminator prediction. All programs were used to obtain genome-wide terminator predictions. As the experimentally identified terminators in BacTermData are not exhaustive; i.e., there might be actual terminators missing from BacTermData. Predictive performance was evaluated using recall at ten sequence overlapping thresholds between the predicted terminators and the actual terminators (Table 8). To do this, we used BEDtools’ intersect command and calculated recall at each overlap threshold.

**Table 8:** Average recall over ten overlap thresholds ± standard deviation for four other terminator prediction tools and BacTermFinder. The corresponding genome accession and GC content (%) is provided below each bacterium name. The highest recall per row is highlighted in bold.

| Bacterium | RhoTermPredict | ITerm-PseKNC | TransTermHP | TermNN | BacTermFinder |
| --- | --- | --- | --- | --- | --- |
| *Streptomyces gardneri* ATCC 15439<br>CP059991.1, GC=70.73% | 0.2499 ± 0.1263 | 0.0101 ± 0.0042 | 0.1709 ± 0.1035 | 0.3690 ± 0.2227 | **0.6029 ± 0.2177** |
| *Mycobacterium tuberculosis* H37Rv<br>AL123456.3, GC=64.69% | **0.2424 ± 0.1640** | 0.1834 ± 0.1149 | 0.0033 ± 0.0018 | 0.1396 ± 0.0927 | 0.2306 ± 0.1191 |
| *Synechocystis* sp. PCC 7338<br>CP054306.1, GC=47.14% | 0.1040 ± 0.0443 | 0.1178 ± 0.0523 | 0.2786 ± 0.1488 | 0.5136 ± 0.2643 | **0.6361 ± 0.2206** |
| *Synechocystis* sp. PCC 6803<br>NC_000911.1, GC=47.04% | 0.1210 ± 0.0789 | 0.1012 ± 0.0544 | 0.2226 ± 0.1035 | 0.4817 ± 0.2427 | **0.5450 ± 0.1818** |
| *Streptococcus agalactiae* NEM316<br>NC_004368.1, GC=35.12% | 0.0188 ± 0.0062 | 0.2402 ± 0.1256 | 0.5596 ± 0.2656 | 0.7715 ± 0.3196 | **0.8209 ± 0.3002** |
| **Overall mean recall** | 0.15 ± 0.10 | 0.13 ± 0.09 | 0.25 ± 0.20 | 0.46 ± 0.23 | **0.57 ± 0.21** |

We included in the comparative assessment bacteria with different GC content. Bacteria with high GC content tend to have a larger proportion of factor-dependent terminators [17]. For example, *M. tuberculosis* is reported to have a large proportion (up to 54%) of factor-dependent terminators [17].

As RhoTermPredict was designed to predict factor-dependent terminators, we hypothesized it would perform better on bacteria with high GC content than on bacteria with low GC content. On the other hand, we hypothesized that termNN and TransTermHP would perform better on bacteria with low GC content than on bacteria with high GC content, as these two tools were designed to identify intrinsic terminators. Our results (Table 8) indeed support these hypotheses. RhoTermPredict achieved its highest recall (roughly 24%) on *M. tuberculosis* and *S. gardneri* which are the two organisms with the highest proportion of factor-dependent terminators. BacTermFinder’s recall on these two organisms (i.e., *M. tuberculosis* and *S. gardneri*) is comparable to or better than that of RhoTermPredict. All tools but RhoTermPredict achieved their highest recall on *S. agalactiae* (lowest GC content). Tools’ recall varied between species with an overall recall range of [0.03, 0.82]. BacTermFinder outperformed ITerm-PseKNC, TermNN and TransTermHP in terms of mean recall across various overlap thresholds in all the species in the validation set, and it achieved the highest mean recall overall (Table 8). This result suggests that BacTermFinder is able to find both types of terminators (intrinsic and factor-dependent) and can generalize to phyla not seen during training (e.g., Cyanobacteriota).

As we have an incomplete annotation of all terminators in any given bacterial genome, it is hard to estimate the genome-wide false positive rate of terminator prediction tools since a prediction might indeed be correct even though a terminator might not have yet been determined experimentally in that location. However, the number of predicted terminators per gene can provide an estimate of the number of false positives. Figure 9 shows the distribution of number of terminators predicted per gene by TermNN, RhoTermPredict and BacTermFinder across the five bacteria in our independent validation data. BacTermFinder displays less variation in the number of predicted terminators per gene, and predicts, on average, 6.62 ± 1.18 terminators per gene; while TermNN and RhoTermPredict predict 8.89 ± 3.52 and 4.30 ± 3.20, respectively. RhoTermPredict predicts less terminators per gene; however, its recall is substantially lower than BacTermFinder’s for four out of five bacteria in our independent validation data. The results provided in Figure 9 and Table 8 indicate that BacTermFinder’s false positive rate is lower than that of TermNN (the 2nd best tool) while achieving a higher recall rate.

**Figure 9:**
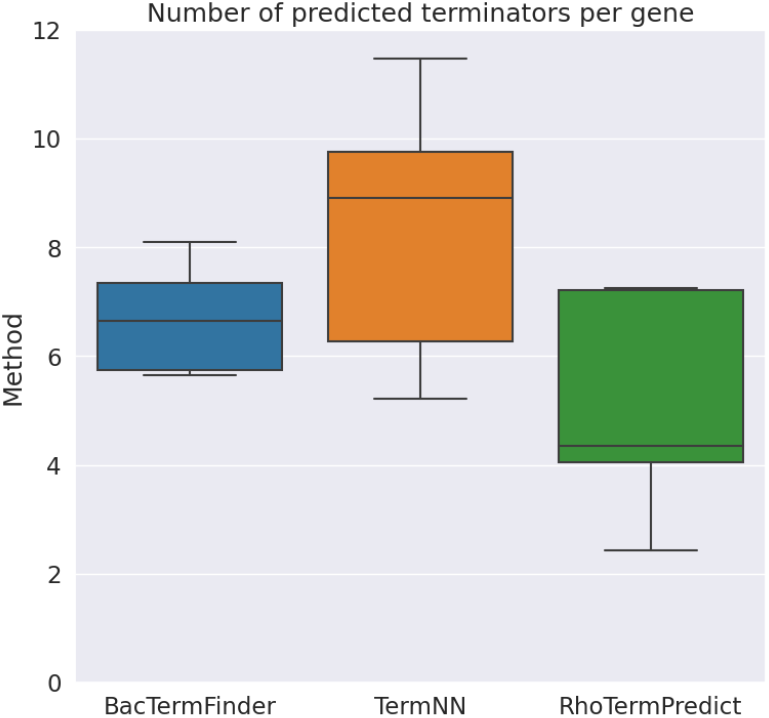
Distribution of the number of predicted terminators per gene across the five bacterial species in our independent validation data. The horizontal line inside each box indicates the median value, and the bottom and top of each box indicate the 25 and 75 percentile, respectively

We used TermNN and BacTermFinder, which are the two tools with the highest recall as per Table 8, to find archaeal terminators. BacTermFinder out-performs TermNN in terms of recall in predicting archaeal terminators (Table 9). This indicates that BacTermFinder can generalize to identify archaeal terminators.

**Table 9:** Average recall over ten overlap thresholds on two archaeal species of the two best performing terminator prediction tools (TermNN and Bac-TermFinder) as per the results shown in Table 8. The highest recall per Archaea is highlighted in bold.

| Archaea | TermNN | BacTermFinder |
| --- | --- | --- |
| *Haloferax volcanii*<br>NC_013967, GC=65.69% | 0.1536 ± 0.1067 | **0.3158 ± 0.1969** |
| *Methanococcus maripaludis*<br>NC_00579, GC=32.63% | 0.4000 ± 0.2235 | **0.5712 ± 0.2468** |

#### 3.5.1 BacTermFinder can predict the location of terminators accurately

We assessed how closely our predictions aligned with actual terminator locations. To do this, we visualized the recall rate as a function of sequence overlap of the predicted terminators with the actual terminators (Fig. 10). We hypothesized that we would observe a declining trend as we moved towards stricter overlap thresholds, which is indeed the case. We compared our overlap vs recall with that of TermNN. Our results show that 1) on every overlap threshold, BacTermFinder outperforms or is comparable to TermNN, and 2) BacTermFinder’s recall drops at stricter overlaps than TermNN’s recall. The latter indicates that BacTermFinder can find the location of terminators more accurately than TermNN. However, BacTermFinder’s recall sharply decreases at overlap thresholds higher than 0.8, which suggests the potential need for a nucleotide-wise segmentation approach to achieve accuracy at the nucleotide level.

**Figure 10:**
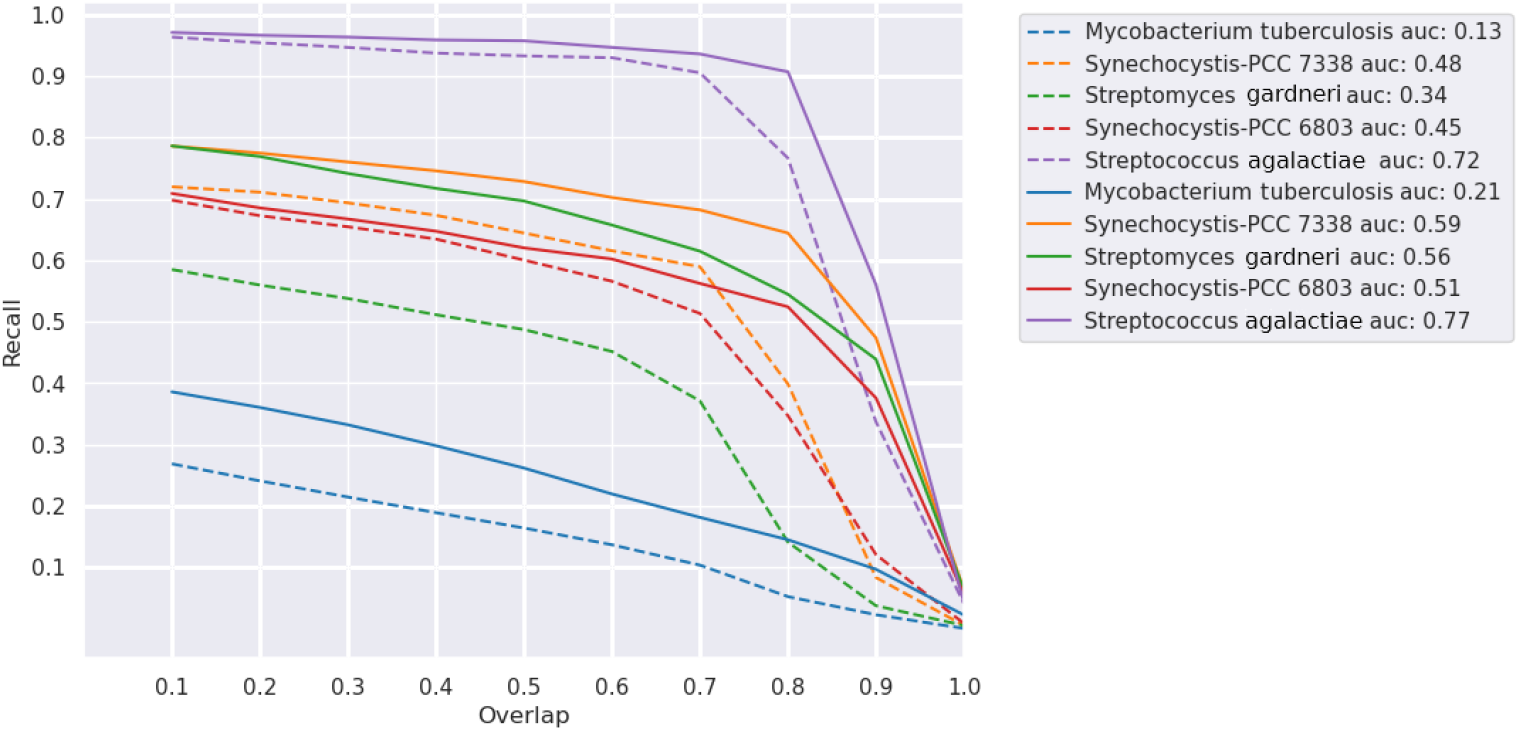
Recall as a function of percentage sequence overlap between predicted and actual terminators. All sequences are 100 nts long. The dotted lines are TermNN, and the solid lines are BacTermFinder. Each colour represents a bacterium in our independent validation data. The area under the curves is shown in the legend.

Additionally, we assessed the accuracy of the probability of being a terminator outputted by TermNN and BacTermFinder. To do this, we sorted their predictions based on their estimated probabilities of a sequence being a terminator, and selected the top *n*% to calculate recall at ten different percentage overlaps with actual terminators. Subsequently, we calculated the mean recall and standard deviation across the percentage overlap levels. We visualized mean recall vs top *n* predictions for a bacterium with high GC content (*S. gardneri*, Fig. 11(A)), a bacterium with approximately equiprobable nucleotide distribution (*Synechocystis* PCC 7338, Fig. 11(B)), and a bacterium with low GC content (*S. gardneri*, Fig. 11(C)). For these three bacteria, BacTermFinder achieves higher recall rates at any top *n*% predictions than TermNN. BacTermFinder has a wider margin over TermNN as the GC content increases. BacTermFinder has also less variation in recall across different overlap thresholds (shaded area in Fig. 11) than TermNN. For *S. gardneri*, the worst recall level of BacTermFinder is similar to or higher than the best recall level of TermNN across all top *n*% of predictions. With BacTermFinder’s 10% most confident predictions on any of these three bacteria, one is able to identify close to 40% of their known terminators.

**Figure 11:**
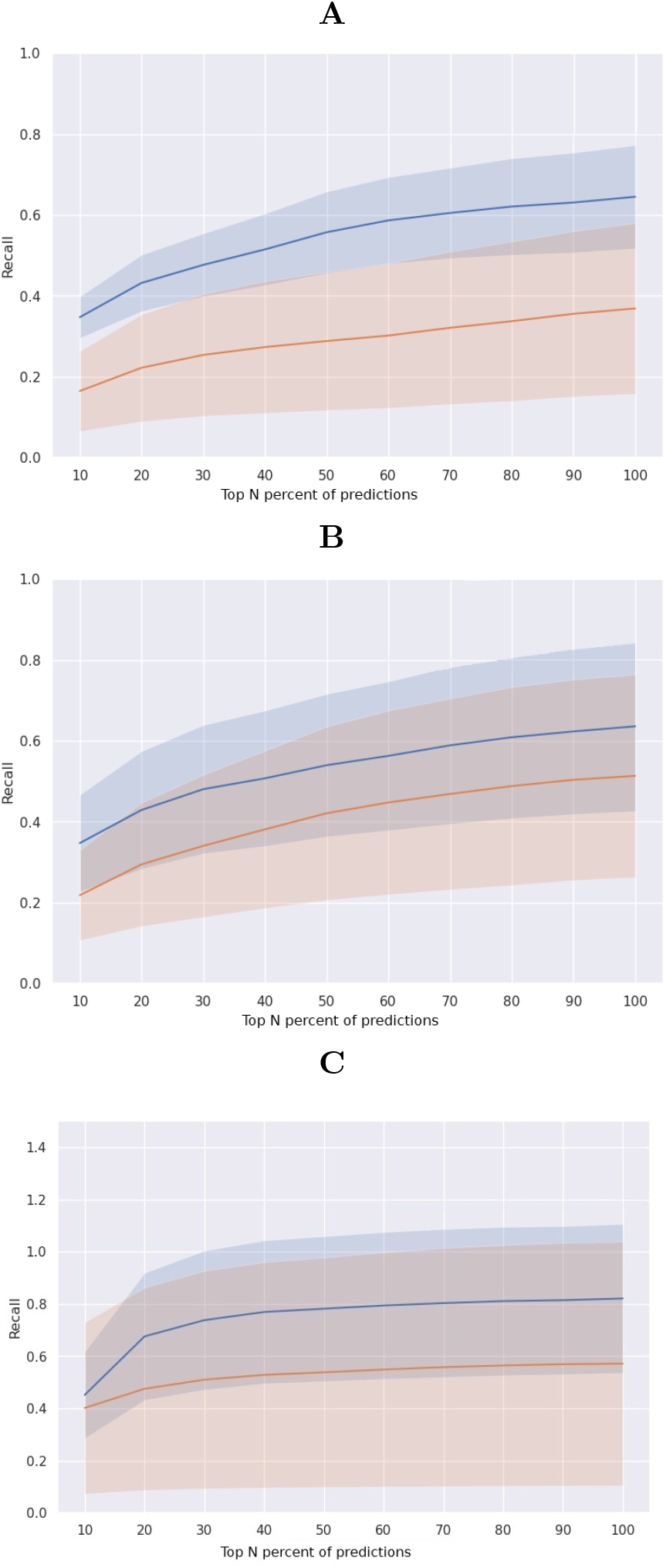
Average recall (solid line) versus top *n*% most confident predictions for BacTermFinder (blue) and TernNN (orange) for **A** *S. gardneri*, **B** *Synechocystis* PCC 7338, and **C** *S. agalactiae*. The shaded area indicates standard deviation.

#### 3.5.2 Low agreement between terminator prediction tools

To see the agreement among RhoTermPredict, TermNN and BacTermFinder, we generated a Venn diagram for two bacteria in the extremes of the GC content axis in our validation data. That is *M. tuberculosis* with 64.69% GC content and *S. agalactiae* with 35.12% GC content. We chose *M. tuberculosis* as it has a large proportion of factor-dependent terminators [17]. For *M. tuberculosis*, the highest agreement (14% of the total predicted terminators) is between BacTermFinder and RhoTermPredict (Fig. 12, yellow and gray areas on the center). For *S. agalactiae*, a low GC bacteria, the highest agreement (92.8% of the total predicted terminators) is between BacTermFinder and TermNN (Fig. 13, pink and gray areas on the left side). This suggests that there is lower agreement among tools to predict factor-dependent terminators (Fig. 12) than intrinsic terminators (Fig. 13). Bac-TermFinder predicts a similar number of terminators in *M. tuberculosis* and *S. agalactiae* as RhoTermPredict and TermNN, respectively. This suggests that BacTermFinder is as effective in identifying factor-dependent and intrinsic terminators as tools specialized on each terminator type. The number of terminators predicted by all three tools is quite low for both bacteria: 23 (or 1.9%) and 9 (or 1.4%) for *M. tuberculosis* and *S. agalactiae*, respectively.

**Figure 12:**
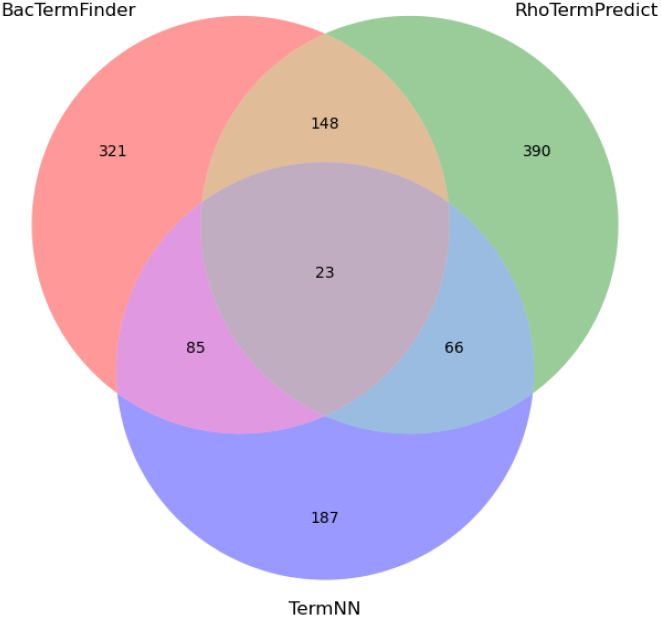
Agreement among 1,220 predicted terminators (with at least 50% sequence overlap with an actual terminator) of BacTermFinder, RhoTermPredict and TermNN for *M. tuberculosis*.

**Figure 13:**
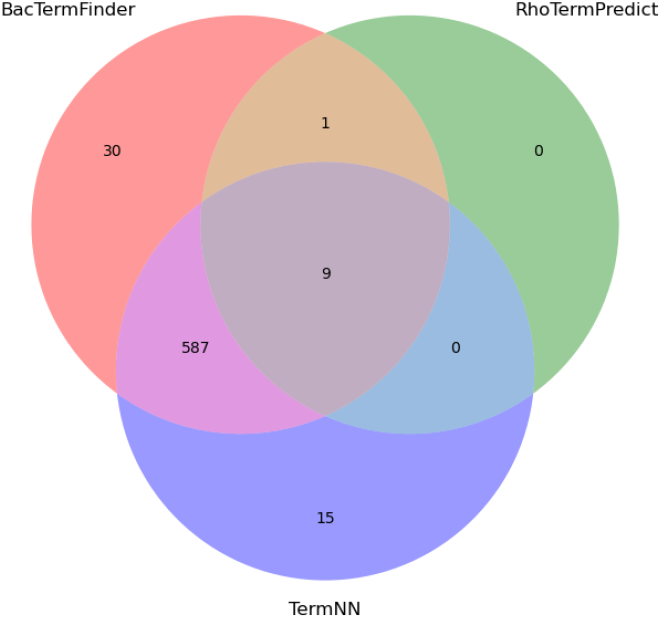
Agreement among 642 predicted terminators (with at least 50% sequence overlap with an actual terminator) of BacTermFinder, RhoTermPredict and TermNN for *S*.*agalactiae*.

## 4 Conclusions

In this work, we have collected, to our knowledge, the largest data set of bacterial terminators identified by sequencing technologies. This comprehensive dataset includes terminators from 25 bacterial strains with a wide range of GC content identified by various sequencing technologies. We expect these data to be valuable for further developments in bacterial terminator prediction. Additionally, we have developed BacTermFinder, a general model for finding bacterial terminators. Future work to improve BacTermFinder includes 1) increasing the region of interest to include the Rho utilization site (rut); 2) adding archaeal terminators into the training data; and 3) determining the terminator type, as BacTermFinder can find both terminator types but does not indicate which type is found.

BacTermFinder outperforms in terms of recall other existing tools in an independent validation data set. BacTermFinder’s average recall over five bacterial species is 0.57 ± 0.21, and TermNN’s (the second-best tool) recall is 0.46 ± 0.23. This increase in recall is achieved by BacTermFinder while predicting, on average, two terminators less per gene than TermNN. BacTermFinder recall in all prokaryotic data (bacterial and archaeal) is 0.53 ± 0.20, and TermNN’s is 0.40 ± 0.22. Furthermore, BacTermFinder can identify both types of terminators as good as or better than tools specialized on that specific terminator type. BacTermFinder and BacTermData are available at: https://github.com/BioinformaticsLabAtMUN/BacTermFinder.

## 6 Author contributions statement

L.P-C. conceived the project, supervised it, interpreted the results, wrote and reviewed the manuscript. S.M.A.T.G. collected BacTermData, implemented BacTermFinder, conducted the experiments, analysed and interpreted the results, wrote and reviewed the manuscript.

## 7 Acknowledgments

This research was partly enabled by computing infrastructure provided by Acenet (ace-net.ca) and the Digital Research Alliance of Canada (alliancecan.ca). This work was supported by funds from an NSERC Discovery Grant to L.P-C. and a graduate fellowship from Memorial University School of Graduate Studies to S.M.A.T.G.

