## Supplementary File for "BacTermFinder: A Comprehensive and General Bacterial Terminator Finder using a CNN Ensemble"

### BacTermFinder's additional file

Seyed Mohammad Amin Taheri Ghahfarokhiand and
Lourdes Peña Castillo
Memorial University of Newfoundland

July 5, 2024

#### Contents

**1 Studies used as data sources for training** **2**

**2 Description of the 1694 features** **3**

**3 Supplementary figures** **4**

**Bibliography** **8**

### 1 Studies used as data sources for training

Table 1: PubMed Central (PMC) identifiers, data accession numbers and sequencing technology per study used during data collection.

| Study name | PMC ID or DOI | GEO or SRA or ENA project # | Sequencing Tech. |
|---|---|---|---|
| [1] | PMC5756622 | PRJEB12568 | Term-Seq |
| [2] | 10.1038/nmicrobiol.2016.143 | PRJEB12568 | Term-Seq |
| [3] | PMC8780764 | GSE118597 | Term-Seq |
| [4] | PMC6742748 | PRJEB31507 | Term-Seq |
| [5] | PMC8269248 | GSE138325 | Term-Seq |
| [6] | PMC6814526 | GSE117737 | Send-Seq |
| [7] | PMC6131387 | GSE117273 | SMRT-Cappable-seq |
| [8] | PMC7815308 | PRJNA640168 | Term-Seq |
| [9] | PMC9023263 | PRJEB36932 | Term-Seq |
| [10] | PMC9226507 | GSE67058 | Term-Seq |
| [11] | PMC8060035 | GSE154522 | Term-Seq |
| [12] | PMC7483943 | GSE53767 GSE95211 GSE108295 | Rend-seq |
| [13] | PMC8914203 | PRJEB40918 | Term-Seq |
| [14] | PMC7566282 | GSE139939 | Term-Seq |
| [15] | PMC7738537 | PRJEB40918 PRJEB31507 PRJEB36379 SRX6937123 SRX6937124 PRJEB36379 | Term-Seq |
| [16] | PMC6786874 | PRJEB31965 | Term-Seq |
| [17] | PMC9217812 | GSE158830 | Term-Seq |
| [18] | PMC6296669 | SRP136114 | Term-Seq |
| [19] | PMC5978003 | GSE95211 | Rend-seq |
| [20] | PMC8237595 | GSE155167 | RNAtag-Seq |
| [21] | PMC8745188 | Available upon Request | RNA-Seq |
| [22] | PMC6212727 | SRP063763 | SMRT-Cappable-seq |
| [23] | PMC6061677 | GSE109766 | Term-Seq |
| [24] | PMC8577284 | prjna775855 | Direct RNA-seq |

### 2 Description of the 1694 features

Table 2: Table of selected important features based on SHAP and Feature Importance of LightGBM

| Feature set | Descriptor group | # of important features | Description |
|---|---|---|---|
| Geary | Autocorrelation | 255 | The distribution of amino acid properties throughout the sequence [25]. |
| NMBroto | Autocorrelation | 255 | The distribution of amino acid properties throughout the sequence [26]. |
| ENAC | Nucleic acid composition | 251 | Computes the frequency of each nucleic acid type using a sliding sequence window [27]. |
| PseKNC | Pseudo nucleic acid composition | 181 | K-tuple nucleotide composition [28, 29]. |
| SCPseDNC | Pseudo nucleic acid composition | 104 | Series correlation pseudo dinucleotide composition information [28, 29]. |
| TACC | Autocorrelation and cross-covariance | 100 | The relationship between either the same or different physicochemical indices for trinucleotides separated by a lag distance along the sequence [28]. |
| SCPseTNC | Pseudo nucleic acid composition | 70 | Correlation pseudo trinucleotide composition [28, 29]. |
| PCPseTNC | Pseudo nucleic acid composition | 60 | Considers parallel correlation pseudo trinucleotide composition information [28, 29]. |
| DACC | Autocorrelation and cross-covariance | 57 | The relationship between either the same or different physicochemical indices for dinucleotides separated by a lag distance along the sequence [28, 30, 31]. |
| Z_curve_144bit | Nucleic acid composition | 56 | Frequencies of phase-specific tri-nucleotides [32]. |
| Mismatch | Nucleic acid composition | 43 | The occurrence of kmers, allowing at most m mismatches [33]. |
| PS2 | Residue composition | 41 | Encoded 16 pairs of adjacent pairwise nucleotides (dinucleotides) [33, 34]. |
| CKSNAP | Nucleic acid composition | 39 | K-spaced nucleic acid pairs [27]. |
| NCP | Nucleic acid composition | 35 | Nucleotide chemical property [35]. |
| ANF | Nucleic acid composition | 28 | Accumulated nucleotide frequency [35]. |
| MMI | Mutual information | 26 | Multivariate mutual information [36]. |
| binary | Residue composition | 22 | Each amino acid is represented by a 4-dimensional binary vector [37, 38]. |
| RCKmer | Nucleic acid composition | 19 | Reverse complement kmer [39, 40]. |
| Z_curve_48bit | Nucleic acid composition | 18 | Frequencies of phase independent tri-nucleotides [32]. |
| PCPseDNC | Pseudo nucleic acid composition | 17 | Parallel correlation pseudo dinucleotide composition [41, 29]. |
| Z_curve_9bit | Nucleic acid composition | 9 | Phase-specific mononucleotides [32]. |
| LPDF | Nucleic acid composition | 8 | Local position-specific dinucleotide frequency [42]. |

### 3 Supplementary figures

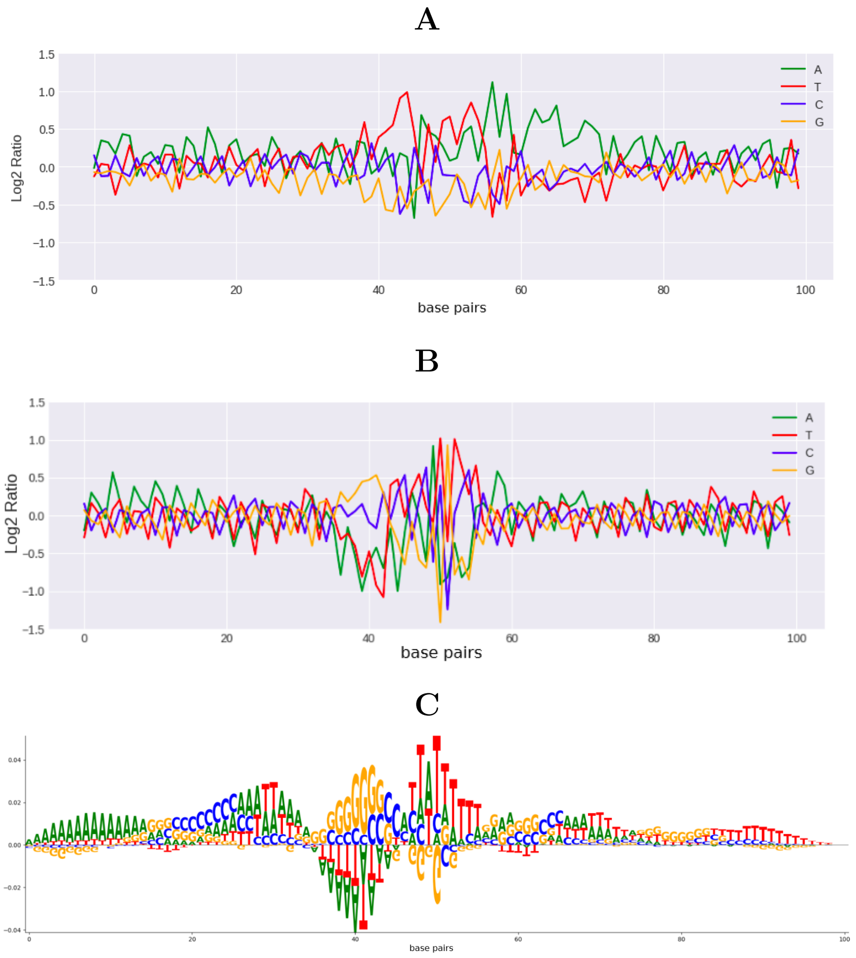


Figure 1: Visualization of *H. volcanii* experimentally identified terminators (A) vs BacTermFinder genome-wide predictions (B). **A**. Log2 ratio of the relative nucleotide frequency of experimentally determined terminators. **B**. Log2 ratio of the relative nucleotide frequency of genome-wide predicted terminators. **C**. Saliency map generated using using [43].

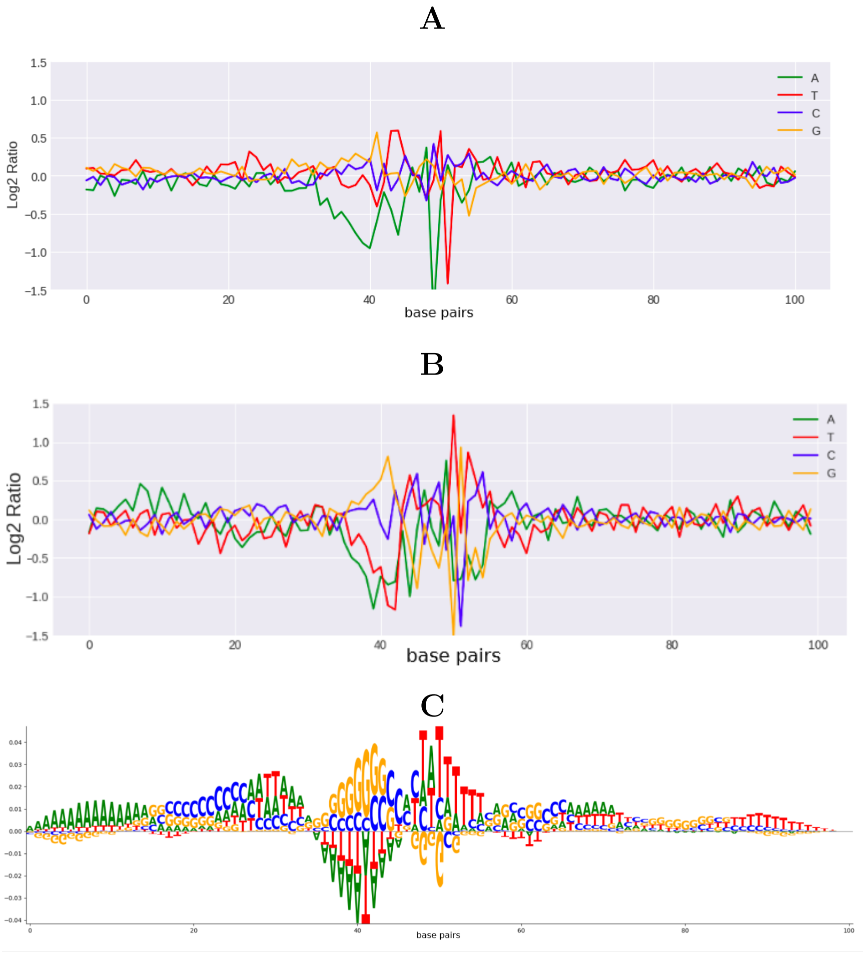


Figure 2: Visualization of *M. tuberculosis* experimentally identified terminators (A) vs BacTermFinder genome-wide predictions (B). **A**. Log2 ratio of the relative nucleotide frequency of experimentally determined terminators. **B**. Log2 ratio of the relative nucleotide frequency of genome-wide predicted terminators. **C**. Saliency map generated using using [43].

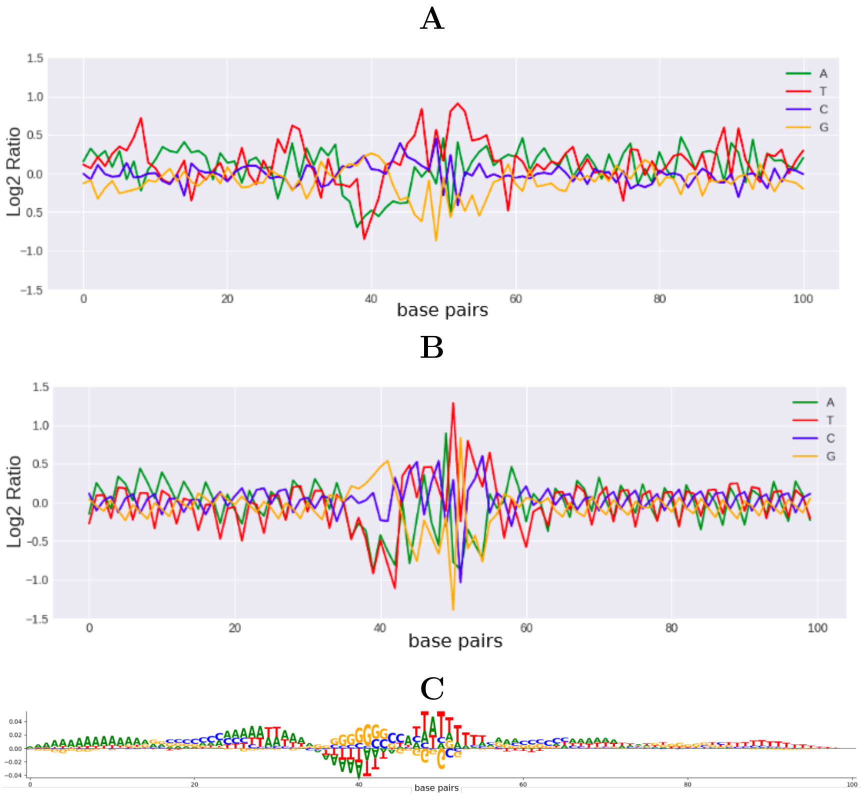


Figure 3: Visualization of *S. gardneri* experimentally identified terminators (A) vs BacTermFinder genome-wide predictions (B). **A**. Log2 ratio of the relative nucleotide frequency of experimentally determined terminators. **B**. Log2 ratio of the relative nucleotide frequency of genome-wide predicted terminators. **C**. Saliency map generated using using [43].

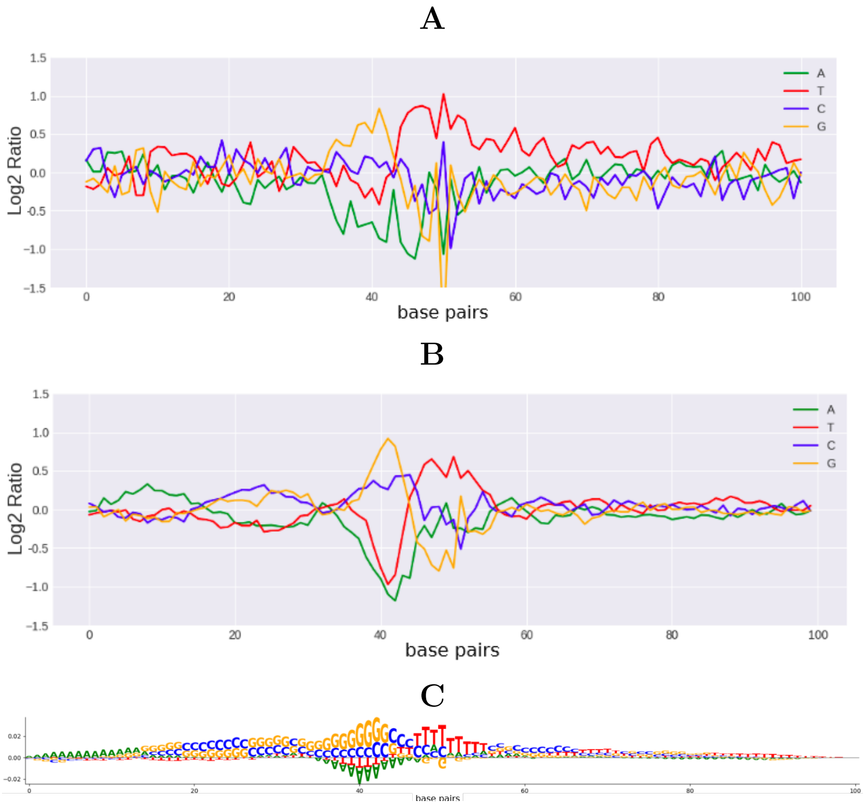


Figure 4: Visualization of *Synechocystis* PCC 7338 experimentally identified terminators (A) vs BacTermFinder genome-wide predictions (B). **A**. Log2 ratio of the relative nucleotide frequency of experimentally determined terminators. **B**. Log2 ratio of the relative nucleotide frequency of genome-wide predicted terminators. **C**. Saliency map generated using using [43].

[3] Yongjae Lee, Namil Lee, Soonkyu Hwang, Woori Kim, Suhyung Cho, Bernhard O. Palsson, and Byung Kwan Cho. Genome-scale analysis of genetic regulatory elements in *Streptomyces avermitilis MA-4680* using transcript boundary information. *BMC Genomics*, 23, 12 2022.

[4] Yongjae Lee, Namil Lee, Yujin Jeong, Soonkyu Hwang, Woori Kim, Suhyung Cho, Bernhard O. Palsson, and Byung Kwan Cho. The transcription unit architecture of *Streptomyces lividans TK24*. *Frontiers in Microbiology*, 10:2074, 9 2019.

[5] Soonkyu Hwang, Namil Lee, Donghui Choe, Yongjae Lee, Woori Kim, Yujin Jeong, Suhyung Cho, Bernhard O. Palsson, and Byung-Kwan Cho. Elucidating the regulatory elements for transcription termination and posttranscriptional processing in the *Streptomyces clavuligerus* genome. *mSystems*, 6, 6 2021.

[6] Xiangwu Ju, Dayi Li, and Shixin Liu. Full-length RNA profiling reveals pervasive bidirectional transcription terminators in bacteria. *Nature microbiology*, 4:1907, 11 2019.

[7] Bo Yan, Matthew Boitano, Tyson A. Clark, and Laurence Ettwiller. SMRT-Cappable-seq reveals complex operon variants in bacteria. *Nature Communications*, 9, 12 2018.

[8] Philip P. Adams, Gabriele Baniulyte, Caroline Esnault, Kavya Chegireddy, Navjot Singh, Molly Monge, Ryan K. Dale, Gisela Storz, and Joseph T. Wade. Regulatory roles of *Escherichia coli* 5' UTR and ORF-internal RNAs detected by 3' end mapping. *eLife*, 10:1–33, 1 2021.

[9] Donghui Choe, Kangsan Kim, Minjeong Kang, Seung Goo Lee, Suhyung Cho, Bernhard Palsson, and Byung Kwan Cho. Synthetic 3'-UTR valves for optimal metabolic flux control in *Escherichia coli*. *Nucleic Acids Research*, 50:4171, 4 2022.

[10] Hiraku Takada, Zachary F. Mandell, Helen Yakhnin, Anastasiya Glazyrina, Shinobu Chiba, Tatsuaki Kurata, Kelvin J.Y. Wu, Ben I.C. Tresco, Andrew G. Myers, Gemma C. Aktinson, Paul Babitzke, and Vasili Hauryliuk. Expression of *Bacillus subtilis* ABCF antibiotic resistance factor vmlr is regulated by RNA polymerase pausing, transcription attenuation, translation attenuation and (p)ppgpp. *Nucleic Acids Research*, 50:6174, 6 2022.

[11] Zachary F. Mandell, Reid T. Oshiro, Alexander V. Yakhnin, Rishi Vishwakarma, Mikhail Kashlev, Daniel B. Kearns, and Paul Babitzke. NusG is an intrinsic transcription termination factor that stimulates motility and coordinates gene expression with NusA. *eLife*, 10, 2021.

[12] Grace E. Johnson, Jean Benoît Lalanne, Michelle L. Peters, and Gene Wei Li. Functionally uncoupled transcription-translation in *Bacillus subtilis*. *Nature*, 585:124, 9 2020.

[13] Soonkyu Hwang, Namil Lee, Donghui Choe, Yongjae Lee, Woori Kim, Ji Hun Kim, Gahyeon Kim, Hyeseong Kim, Neung Ho Ahn, Byoung Hee Lee, Bernhard O. Palsson, and Byung Kwan Cho. System-level analysis of transcriptional and translational regulatory elements in *Streptomyces griseus*. *Frontiers in Bioengineering and Biotechnology*, 10, 2 2022.

[14] Jessica M. Vera, Indro Neil Ghosh, Yaoping Zhang, Alex S. Hebert, Joshua J. Coon, and Robert Landick. Genome-scale transcription-translation mapping reveals features of *Zymomonas mobilis* transcription units and promoters. *mSystems*, 5, 8 2020.

[15] Yongjae Lee, Namil Lee, Soonkyu Hwang, Woori Kim, Yujin Jeong, Suhyung Cho, Bernhard O. Palsson, and Byung Kwan Cho. Genome-scale determination of 5´ and 3´ boundaries of RNA transcripts in *Streptomyces genomes*. *Scientific Data*, 7, 12 2020.

[16] Maureen K. Thomason, Maya Voichek, Daniel Dar, Victoria Addis, David Fitzgerald, Susan Gottesman, Rotem Sorek, and E. Peter Green-

berga. A rhli 5' UTR-derived sRNA regulates RhlR-dependent quorum sensing in *Pseudomonas aeruginosa. mBio*, 10, 2019.

[17] Daniel G. Mediati, Julia L. Wong, Wei Gao, Stuart McKellar, Chi Nam Ignatius Pang, Sylvania Wu, Winton Wu, Brandon Sy, Ian R. Monk, Joanna M. Biazik, Marc R. Wilkins, Benjamin P. Howden, Timothy P. Stinear, Sander Granneman, and Jai J. Tree. RNase III-CLASH of multi-drug resistant *Staphylococcus aureus* reveals a regulatory mRNA 3'UTR required for intermediate vancomycin resistance. *Nature Communications*, 13, 12 2022.

[18] Indu Warrier, Nikhil Ram-Mohan, Zeyu Zhu, Ariana Hazery, Haley Echlin, Jason Rosch, Michelle M. Meyer, and Tim van Opijnen. The transcriptional landscape of *Streptococcus pneumoniae TIGR4* reveals a complex operon architecture and abundant riboregulation critical for growth and virulence. *PLoS Pathogens*, 14, 12 2018.

[19] Jean Benoît Lalanne, James C. Taggart, Monica S. Guo, Lydia Herzel, Ariel Schieler, and Gene Wei Li. Evolutionary convergence of pathway-specific enzyme expression stoichiometry. *Cell*, 173:749, 4 2018.

[20] Manuela Fuchs, Vanessa Lamm-Schmidt, Johannes Sulzer, Falk Ponath, Laura Jenniches, Joseph A. Kirk, Robert P. Fagan, Lars Barquist, Jörg Vogel, and Franziska Faber. An RNA-centric global view of *Clostridioides difficile* reveals broad activity of Hfq in a clinically important gram-positive bacterium. *Proceedings of the National Academy of Sciences of the United States of America*, 118, 6 2021.

[21] Laurène Bastet, Pilar Bustos-Sanmamed, Arancha Catalan-Moreno, Carlos J. Caballero, Sergio Cuesta, Leticia Matilla-Cuenca, Maite Villanueva, Jaione Valle, Iñigo Lasa, and Alejandro Toledo-Arana. Regulation of heterogenous LexA expression in *Staphylococcus aureus* by an antisense RNA originating from transcriptional read-through upon natural mispairings in the sbrB intrinsic terminator. *International Journal of Molecular Sciences*, 23, 1 2022.

[22] Jelle Slager, Rieza Aprianto, and Jan Willem Veening. Deep genome annotation of the opportunistic human pathogen *Streptococcus pneumoniae D39. Nucleic Acids Research*, 46:9971, 11 2018.

[23] Daniel Dar and Rotem Sorek. High-resolution RNA 3'-ends mapping of bacterial Rho-dependent transcripts. *Nucleic Acids Research*, 46:6797, 7 2018.

[24] Mohamad Al kadi, Eiji Ishii, Dang Tat Truong, Daisuke Motooka, Shigeaki Matsuda, Tetsuya Iida, Toshio Kodama, and Daisuke Okuzaki. Direct RNA sequencing unfolds the complex transcriptome of *Vibrio parahaemolyticus*. *mSystems*, 6, 12 2021.

[35] Wei Chen, Hong Tran, Zhiyong Liang, Hao Lin, and Liqing Zhang. Identification and analysis of the N(6)-methyladenosine in the *Saccharomyces cerevisiae* transcriptome. *Scientific reports*, 5, 9 2015.

[36] Leyi Wei, Ran Su, Shasha Luan, Zhijun Liao, Balachandran Manavalan, Quan Zou, and Xiaolong Shi. Iterative feature representations improve N4-methylcytosine site prediction. *Bioinformatics (Oxford, England)*, 35:4930–4937, 12 2019.

[37] Zhen Chen, Yuan Zhou, Jiangning Song, and Ziding Zhang. hck-saap_ubsite: improved prediction of human ubiquitination sites by exploiting amino acid pattern and properties. *Biochimica et biophysica acta*, 1834:1461–1467, 2013.

[38] Zhen Chen, Yong Zi Chen, Xiao Feng Wang, Chuan Wang, Ren Xiang Yan, and Ziding Zhang. Prediction of ubiquitination sites by using the composition of k-spaced amino acid pairs. *PloS one*, 6, 2011.

[39] William Stafford Noble, Scott Kuehn, Robert Thurman, Man Yu, and John Stamatoyannopoulos. Predicting the in vivo signature of human

gene regulatory sequences. *Bioinformatics (Oxford, England)*, 21 Suppl 1, 2005.

[40] Shobhit Gupta, Jonathan Dennis, Robert E. Thurman, Robert Kingston, John A. Stamatoyannopoulos, and William Stafford Noble. Predicting human nucleosome occupancy from primary sequence. *PLoS computational biology*, 4, 2008.

[41] Bin Liu, Fule Liu, Longyun Fang, Xiaolong Wang, and Kuo Chen Chou. repDNA: a Python package to generate various modes of feature vectors for DNA sequences by incorporating user-defined physicochemical properties and sequence-order effects. *Bioinformatics (Oxford, England)*, 31:1307–1309, 4 2015.

[42] Xiaoli Qiang, Huangrong Chen, Xiucai Ye, Ran Su, and Leyi Wei. M6AMRFS: Robust prediction of N6-methyladenosine sites with sequence-based features in multiple species. *Frontiers in genetics*, 9, 10 2018.

[43] Antonio Majdandzic, Chandana Rajesh, and Peter K. Koo. Correcting gradient-based interpretations of deep neural networks for genomics. *Genome Biology*, 24(1):109, 2023.
